# Single-cell analysis of early antiviral gene expression reveals a determinant of stochastic *IFNB1* expression

**DOI:** 10.1101/182972

**Authors:** Sultan Doğanay, Maurice Youzong Lee, Alina Baum, Jessie Peh, Sun-Young Hwang, Joo-Yeon Yoo, Paul J. Hergenrother, Adolfo García-Sastre, Sua Myong, Taekjip Ha

## Abstract

RIG-I-like receptors (RLRs) are cytoplasmic sensors of viral RNA that trigger the signaling cascade that leads to type I interferon (IFN) production. Transcriptional induction of RLRs by IFN is believed to play the role of positive feedback to further amplify viral sensing. We found that RLRs and several other IFN-stimulated genes (ISGs) are induced early in viral infection independent of IFN. Expression of these early ISGs requires IRF3/IRF7 and is highly correlated amongst them. Simultaneous detection of mRNA of *IFNB1*, viral replicase, and ISGs revealed distinct populations of *IFNB1* expressing and non-expressing cells which are highly correlated with the levels of early ISGs but are uncorrelated with IFN-dependent ISGs and viral gene expression. Individual expression of RLRs made *IFNB1* expression more robust and earlier, suggesting a causal relation between levels of RLR and induction of IFN.

## Introduction

RIG-I-like receptors (RLRs) RIG-I, MDA5, and LGP2 constitute a family of cytoplasmic sensors of viral RNA with indispensable roles in innate immunity ^1-5^. Knockout mice that lack *RIG-I* or *MDA5* are highly susceptible to infection by the respective RNA viruses that these sensors recognize ^3^. Knockdown of *RIG-I* expression in fibroblasts, epithelial cells, conventional dendritic cells and macrophages resulted in failure to activate type I IFN upon infection with various RNA viruses, highlighting the indispensable antiviral role of RLRs in a broad range of cell types ^4^. Type I IFN is known to increase transcription of hundreds of IFN-stimulated genes (ISGs), including RLRs, that can effectively combat the infection. A screening of more than 380 human ISGs for their ability to inhibit the replication of several clinically important human and animal viruses showed that RIG-I and MDA5 are among the top five in their antiviral activities against a broad range of viruses ^6^.

After viral RNA recognition, RIG-I and MDA5 interact with the mitochondrial antiviral signaling protein (MAVS), and through a series of signaling cascades, activate transcription of type I IFN which includes *IFNB1* and *IFNA* subtypes ^7-10^. Type I IFN binds to the IFNα/β receptor (IFNAR) on the cell membrane, and activates transcription of ISGs through JAK-STAT pathway, resulting in an antiviral state that controls infection ^11-13^. Some ISGs such as *Viperin, IFIT1* and *OASL* are not only IFN-inducible, but also induced directly by viral infection in the absence of IFN ^14-16^.

Induction of RLRs by IFN is believed to play the role of positive feedback to further amplify viral sensing, however, their expression kinetics during viral infection, and in particular with respect to type I IFN production, has not been systematically analyzed, especially at the early stages of infection, where highly sensitive methods are needed to capture the small changes in gene expression. Majority of the earlier works are based on bulk measurements, where results are ensemble-averaged over a large number of cells, which may hide changes in gene expression, if these changes originated from a small fraction of the cells.

In fact, it is well recognized that there is significant cell-to-cell heterogeneity in *IFNB1* expression such that only a fraction of cells produces IFN-β upon viral infection^17-24^. Several factors have been suggested to contribute to this heterogeneity including infecting virus quasispecies, complexities of multiple transcription factor binding to the *IFNB1* promoter,^19, 22-26^ cellular heterogeneity in the host gene expression levels and paracrine signaling ^19, 22-26^. For example, components of the RIG-I signaling pathway, such as RIG-I, MDA5 and TRIM25 are expressed at higher levels in the IFN-β expressing cells ^22^, and the fraction of IFN-β producing cells increased when RIG-I signaling pathway components such as RIG-I, TRIM25, NF-kB, IRF3 and IRF7 were overexpressed in cells prior to viral infection ^19, 22^. Shalek *et al* have shown that cellular variation is controlled in a paracrine manner in dendritic cells^26^, while Rand *et al*and Patil *et al* have shown in fibroblasts and dendritic cells that heterogeneity in *IFNB1*expression is predominantly cell-intrinsic at early time points post infection, which is then propagated by paracrine response^23, 24^.

As RLRs are major players in viral recognition and subsequent IFN induction, and have been implicated in the heterogeneous expression of *IFNB1*, single-cell measurements with high precision and sensitivity are needed to more accurately understand their expression kinetics and functional consequences in antiviral signaling. We studied host and viral mRNA expression in single cells using single-molecule fluorescence in situ hybridization (smFISH) ^27, 28^. The single cell approach with the ultimate resolution of single transcripts enabled the discovery of a virus-induced, IFN-independent up-regulation of *RIG-I* that is hidden in ensemble analysis. This IFN - independent mechanism operates as early as 3 hours post infection, requires the IRF3/IRF7 pathway, and induces not only *RIG-I* but also several other ISGs, that we will refer to as “early ISGs”, prior to IFN production.

Multi-color smFISH analysis revealed that the mRNA levels of early ISGs are highly correlated with the large cell-to-cell heterogeneity in *IFNB1* expression, in contrast to the viral gene and IFN-dependent ISG levels that are uncorrelated with *IFNB1* expression. Over expression of RIG - I and MDA5 made *IFNB1* expression more robust and earlier, indicating that early expression of *RIG-I* and *MDA5* in a subset of infected cells may contribute to the decision making process for turning on the paracrine IFN-dependent innate immune response.

## Results

### Quantification of antiviral gene expression with single-cell resolution

We performed smFISH experiments on fixed cells ^27^, where each mRNA is decorated with 38 to 48 different sequences of fluorescently labeled probes, each 20 bases long (Fig. 1*A*). The large number of fluorophores bound to a single mRNA yielded a diffraction-limited fluorescent spot clearly above the background. smFISH assay also revealed the active transcription sites in the nucleus as brighter fluorescent spots where nascent mRNA molecules accumulate. For simultaneous quantification of up to three different genes in single cells, we labeled the probes with spectrally distinguishable fluorophores (Cy3, Alexa Fluor 594 and Cy5) (Fig. 1*A*), and counted the fluorescent spots from the 3D images of cells ^27^.

**Figure 1.**
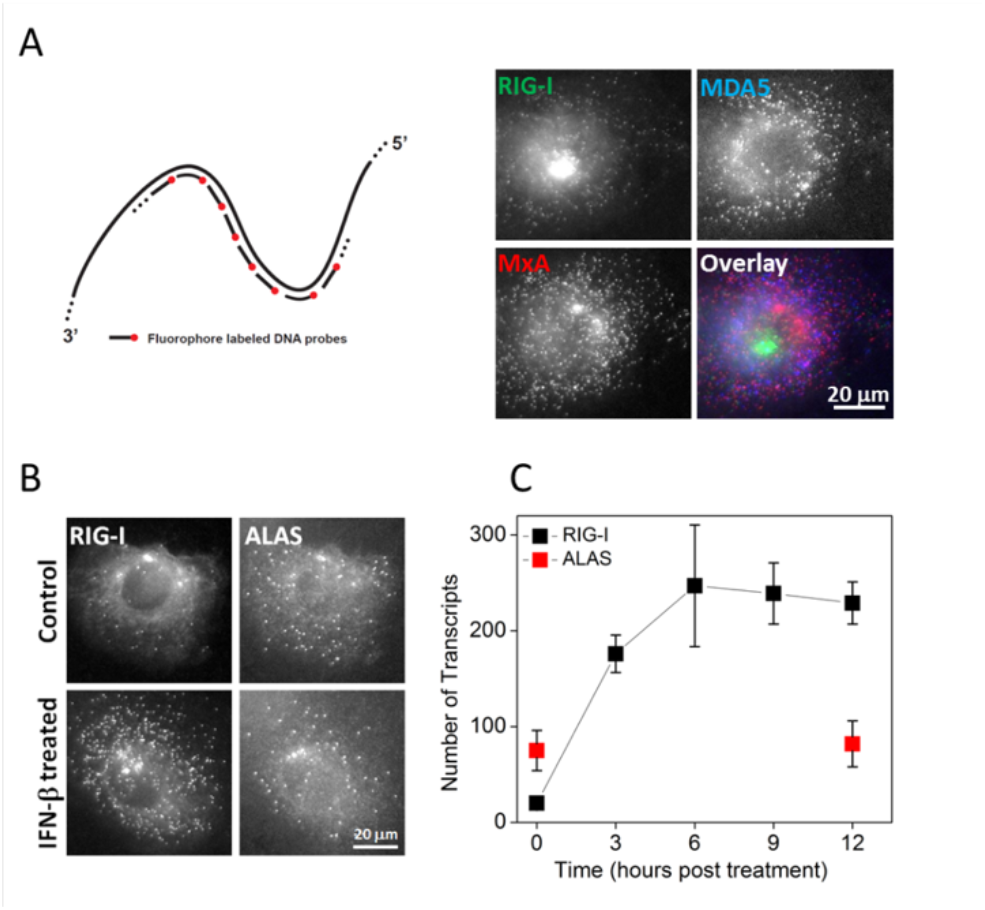
Quantification of IFN induced gene expression in single HepG2 cells (A) The schematics of single-molecule fluorescence in-situ hybridization (smFISH) and simultaneous imaging of *RIG-I, MXA,* and *MDA5* mRNA in individual HepG2 cells treated with IFN-β for 6hours. (B) Images of *RIG-I* and housekeeping gene *ALAS* mRNA in control and IFN-β treated (6 hours) HepG2 cells. (C) Kinetics of *RIG-I* mRNA expression post IFN-β treatment. Transcript numbers (mean ± SD) are obtained from 10 to 20 cells at each time point.

We first examined IFN-induced *RIG-I* expression. We treated previously described clonal HepG2 cells ^29^ with IFN-β and imaged *RIG-I* and housekeeping gene *ALAS* ^30^ mRNA at various time points. Unstimulated cells displayed basal level of *RIG-I* mRNA, 27 transcripts per cell on average. IFN-β treatment changed *RIG-I* mRNA counts in a time-dependent manner with the average number increasing with time but the mean intensity of individual spots remaining constant (Fig. 1*A, B, C*, Fig. S1), supporting the detection of individual transcripts. The cytoplasmic *RIG-I* mRNA counts displayed a steep increase until 6 hours followed by a plateau with an average of >200 transcripts per cell (Fig. 1*C*). In contrast, *ALAS* mRNA counts stayed relatively unchanged.

### *RIG-I* is directly induced at the early stages of viral infection, independent of IFN

To examine the virus-induced expression of *RIG-I* and *IFNB1*, we infected clonal HepG2 cells with Sendai virus (SeV), fixed the cells at various time points post infection, and imaged *RIG-I*and *IFNB1* mRNAs. The earliest time point at which we could detect *IFNB1* transcripts was 9 hours post infection (hpi) (Fig. 2*A*, *B*). Consistently, IFN-β in the culture medium was detected with ELISA from 9 hpi onwards (Fig. 2*C*). On the other hand, bright *RIG-I* transcription sites were visible in the nuclei of a small fraction of the cells as early as 3 hpi, several hours prior to *IFNB1* induction (Fig. 2*A*, Fig. S2). These cells contained 58 cytoplasmic RIG-I mRNA on average, which is about twice as many as basal level of 27 transcripts (Fig. 2D, Table 1). Percentage of cells expressing *RIG-I* mRNA over basal levels reached 32% by 6 hpi with an average of 145 transcripts per cell (Fig. 2*B*, *D*), and further rose to 80% at 9 hpi. Thus, our results show that *RIG-I* is directly induced at the early stages of viral infection in the absence of detectable *IFNB1*.

**Figure 2.**
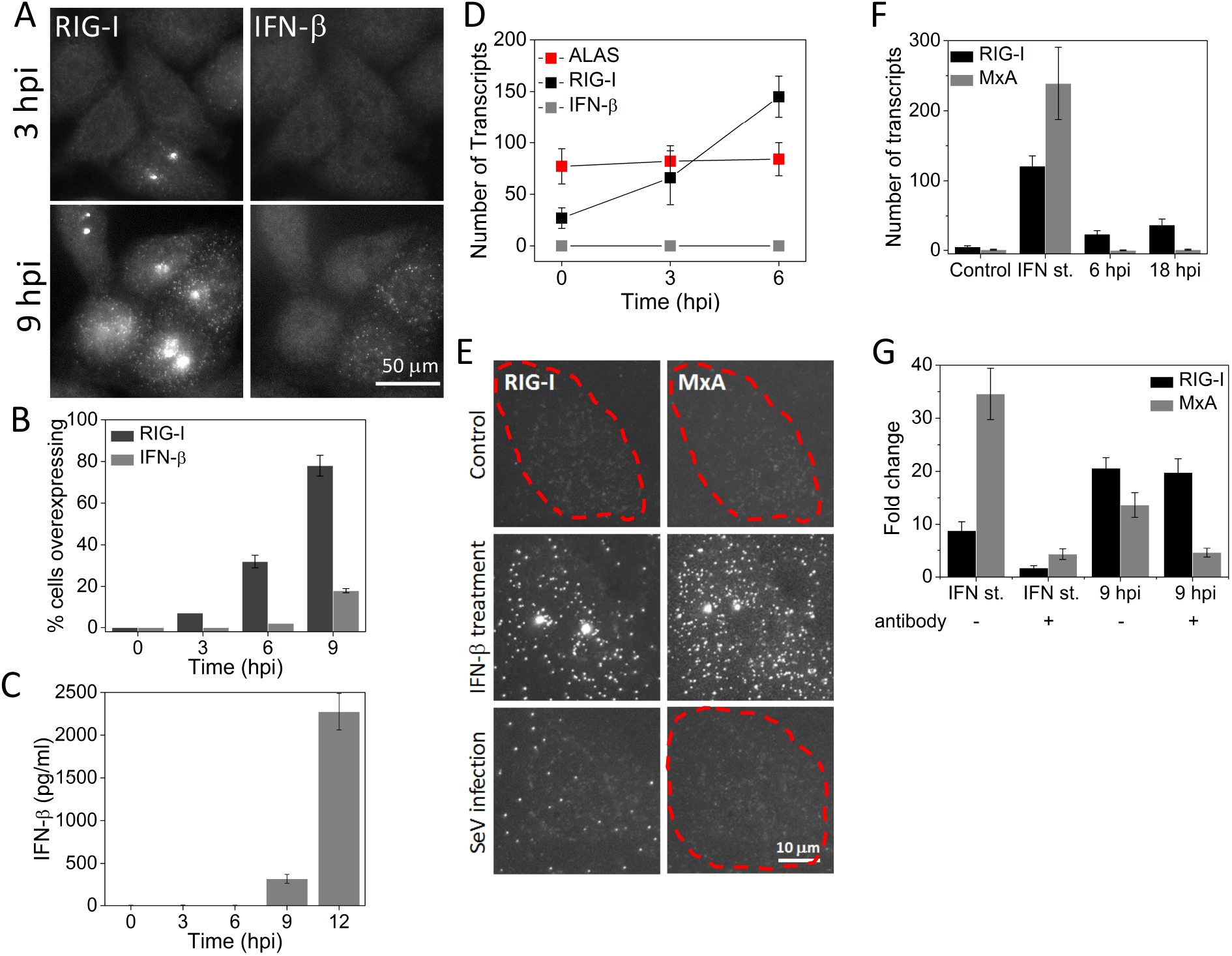
*RIG-I* is induced IFN-independently at the early stages of Sendai virus infection. (A) Images of *RIG-I* and *IFNB1* mRNA in HepG2 cells at 3 and 9 hours post SeV infection. (B) Percentage (mean ± SD) of HepG2 cells overexpressing *RIG-I* and *IFNB1* mRNA as a function of time post SeV infection. 600 to 1000 cells were analyzed at each time point. (C) ELISA to detect IFN-β (mean ± SD) in the culture medium at different time points after SeV infection. Data is obtained from three independent experiments. (D) Number of *RIG-I*, *IFNB1* and *ALAS*transcripts per cell (mean ± SD) in SeV infected (3 and 6 hours) HepG2 cells. Transcript numbers are obtained from 50 cells that overexpress *RIG-I* at each time point. (E) Images of *RIG-I* and *MXA* mRNA in control, IFN-β treated (12 hours) or SeV infected (18 hours) Vero cells. Cell borders are denoted by dashed outlines. (F) Number of *RIG-I* and *MXA* mRNA per cell (mean ± SD) in control, IFN-β treated (12 hours) or SeV infected (6 and 18 hours) Vero cells. 20 cells are analyzed at each condition. (G) qRT-PCR analysis of *RIG-I* and *MXA* mRNA in IFN-β treated (2 hours) or SeV infected (9 hours) HepG2 cells in the presence or absence of neutralizing antibodies against IFN-α, IFN-β, IFN-λ, and IFNAR. Results are normalized to GAPDH, and presented as fold change relative to control cells. Error bars show standard deviation of two independent experiments.

**Table 1.** Number of transcripts (mean ± SD) detected by smFISH in control, IFN-β treated (12 hours), and SeV infected (3 hours) HepG2 cells. Over-expressing cells were chosen to determine average transcript numbers at 3 hpi for genes in the dark grey rows. Cells were chosen randomly to determine the rest of the data. Data obtained from at least 20 cells for each condition and gene. Dark gray rows denote early ISGs.

| | Basal | IFN-β st. | SeV (3 hpi) |
| --- | --- | --- | --- |
| RIG-I | 27 ± 10 | 247 ± 64 | 58 ± 20 |
| MDA5 | 32 ± 12 | 231 ± 60 | 68 ± 18 |
| LGP2 | 7 ± 2 | 44 ± 12 | 31 ± 7 |
| OasL | 0.4 ± 0.5 | 148 ± 44 | 255 ± 52 |
| IFIT1 | 3.5 ± 1.8 | 212 ± 68 | 303 ± 84 |
| MxA | 34 ± 7 | 273 ± 52 | 36 ± 8 |
| NLRX1 | 6 ± 5 | 68 ± 27 | 6 ± 4 |
| TRIM25 | 82 ± 52 | 334 ± 97 | 62 ± 33 |

To confirm that virus-induced early expression of *RIG-I* is indeed independent of IFN, first, we employed Vero cells that cannot produce any type of IFN due to genetic defects, but do have IFNAR and an intact JAK-STAT pathway, and therefore produce ISGs in response to IFN-β stimulation ^31^. As expected, upon IFN-β treatment, Vero cells showed a large increase in *RIG-I*transcription, from less than 5 to ~ 120 transcripts per cell. Upon infection with SeV, we did not detect any *IFNB1* mRNA at any time point while the number of *RIG-I* transcripts gradually increased up to ~50 transcripts per cell, supporting our conclusion that *RIG-I* is induced during viral infection independent of IFN. To exclude the possibility that trace amount of *IFNB1* in the virus stock may give rise to *RIG-I* induction, we also imaged *MXA* mRNA, which is strictly induced by IFN^32^. *MXA* transcript counts remained negligible during infection while externally added *IFNB1* did induce *MXA* up to ~240 transcripts per cell (Fig. 2*E*, *F*).

In another experiment, we used quantitative RT-PCR (qRT-PCR) to determine how *RIG-I*induction is modulated by the neutralizing antibodies against IFN-α, IFN-β, IFN-λ, and IFNAR. In the absence of neutralizing antibodies, *RIG-I* and *MXA* were induced 9 - and 35-fold respectively upon treating the cells with IFN-β (Fig. 2*G*). The addition of the neutralizing antibodies during IFN-β treatment suppressed *RIG-I* and *MXA* induction, showing their efficacy (Fig. 2*G*). At 9 hours post SeV infection, *RIG-I* and *MXA* were induced 21 - and 14-fold respectively in the absence of neutralizing antibodies. Addition of neutralizing antibodies during infection effectively suppressed *MXA* but did not affect *RIG-I* levels (Fig. 2*G*). These results further support that *RIG-I* is mainly regulated by an IFN-independent pathway at the early stages of viral infection, and even at 9 hpi much of *RIG-I* expression occurs independent of the IFN signaling pathway.

### IFN-independent expression of early ISGs is strongly correlated amongst them and is dependent on IRF3/IRF7

Upon further examination using smFISH, expression kinetics of a small subset of ISGs revealed differential regulation of ISGs in response to viral infection. Antiviral genes *RIG-I, MDA5, LGP2*, as well as *IFIT1* and *OASL*, already known to be induced IFN-independently, were directly induced by SeV infection as early as 3 hpi, observed as bright transcription sites in the nucleus and higher cytoplasmic mRNA copy numbers compared to control cells (Table 1). In contrast, *IRF7, NLRX1, MXA, PKR*, and *TRIM25* did not display elevated levels of mRNA at early times post infection, even though they all are induced upon IFN-β treatment (Fig. 3A, B, Table 1) ^13-15^.

**Figure 3.**
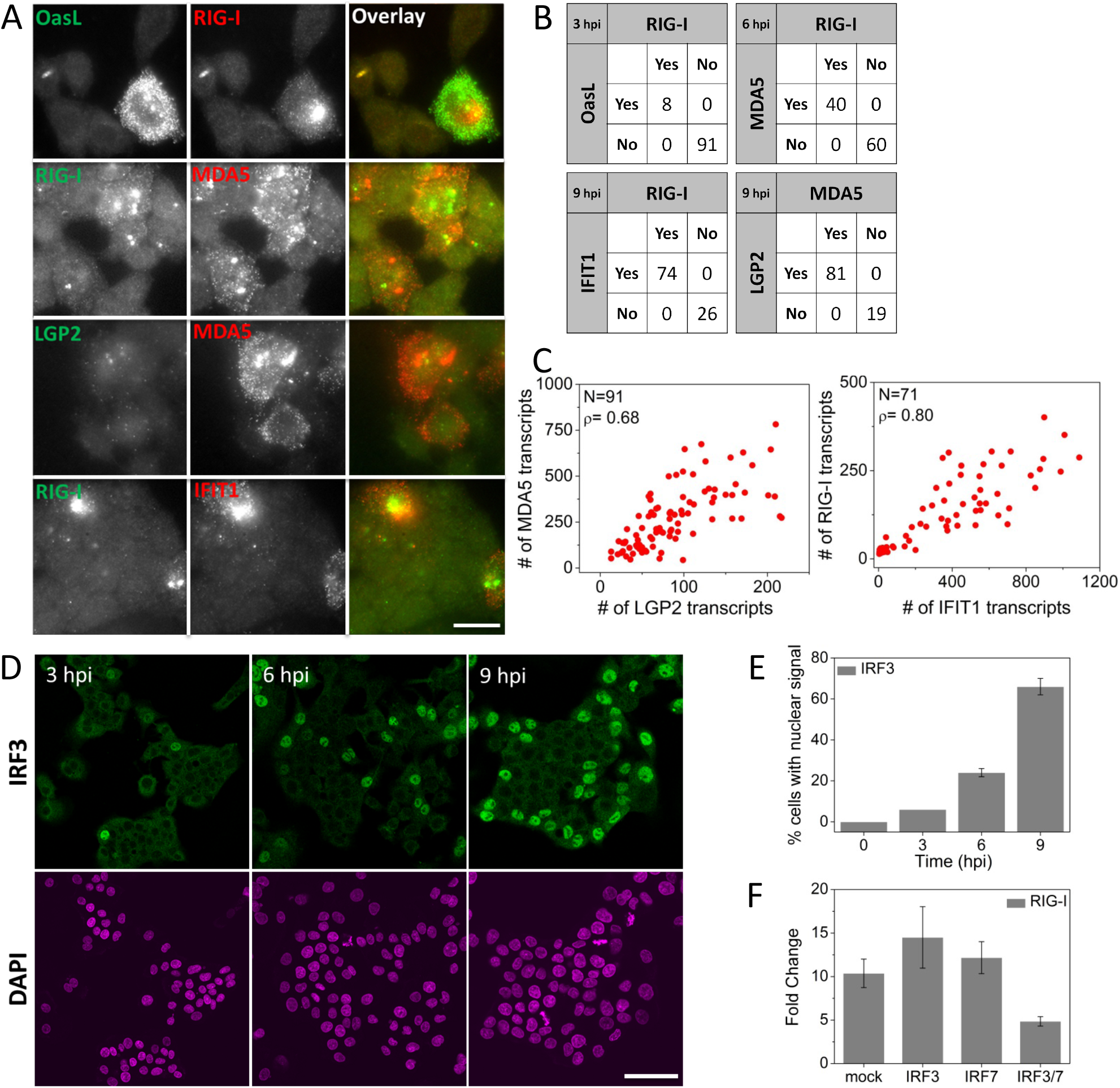
Virus-induced IFN-independent expression of early ISGs is strongly correlated and is dependent on IRF3/IRF7. (A) Pairwise images of *RIG-I* and *OASL*, *RIG-I* and *MDA5*, *LGP2* and *MDA5*, and *RIG-I* and *IFIT1* transcripts in SeV infected HepG2 cells. Scale bar: 50 µm (B) Percentage of cells expressing *RIG-I* and/or *OASL*, *RIG-I* and/or *MDA5*, *MDA5* and/or *LGP2*, and *RIG-I* and/or *IFIT1* at indicated times post SeV infection. 200-300 cells were analyzed for each case. (C) Scatter plots of number of *MDA5* vs. *LGP2*, and *RIG-I* vs. *IFIT1*transcripts in single HepG2 cells at 9 hours post SeV infection. N: Number of cells analyzed, r: Pearson’s correlation coefficient. (D) Images of immunofluorescently labeled IRF3 at various time points post SeV infection. Scale bar: 50 µm (E) Percentage of cells (mean ± SD) displaying nuclear IRF3 as a function of time post SeV infection. 450 to 600 cells were analyzed at each time point. (F) qRT-PCR analysis of *RIG-I* mRNA in SeV infected (9 hours) HepG2 cells with prior siRNA knockdown (30 hours) of mock (HPRT1), IRF3, IRF7 and double knockdown of IRF3 and IRF7. Results are normalized to GAPDH, and presented as fold change relative to control cells. Error bars show standard deviation of two independent experiments.

Two-color smFISH analysis of single cells revealed that SeV-induced expressions of early ISGs are highly correlated with each other. Pairwise analyses of *RIG-I* vs. *MDA5*, *MDA5* vs. *LGP2*, *RIG-I* vs. *OASL* and *RIG-I* vs. *IFIT1* showed that if a cell activates transcription one of these genes, the other paired gene is also activated (Fig. 3*A*, *B*). For example, at 3 hpi, 92% of cells showed no induction of either *RIG-I* or *OASL*, 8% of cells showed both, and none showed *RIG-I*only or *OASL* only. We obtained high Pearson’s correlation coefficients for the mRNA counts for early ISGs in individual cells, for instance 0.8 between *RIG-I* and *IFIT1*, and 0.68 between *LGP2* and *MDA5* 9 hpi (Fig. 3*C*).

Next, we used the Nanostring nCounter system ^33^ and tested a panel of 49 innate immunity genes for their early expression during infection, as well as to compare an ensemble assay to our single cell assay. nCounter data obtained from an ensemble of cells confirmed that first considerable *IFNB1* expression happens at 9 hpi. (Table 2). At 6 hours post SeV infection, we identified 8 out of the 49 genes that are up-regulated (more than 2-fold increase with respect to the uninfected cells) in the absence of *IFNB1*: *DDx60, ISG15, IFIT1, LGP2, MDA5, OASL, RIG-I,* and *Viperin*(Table 2). Several other genes that strongly responded to IFN-β treatment, such as *IFITM3, MXA, Myd88, OAS1, PKR,* and *TRIM25* were not induced (less than 1.5-fold increase with respect to the uninfected cells) at 6 hpi (Table 2, Table S1).

**Table 2.**
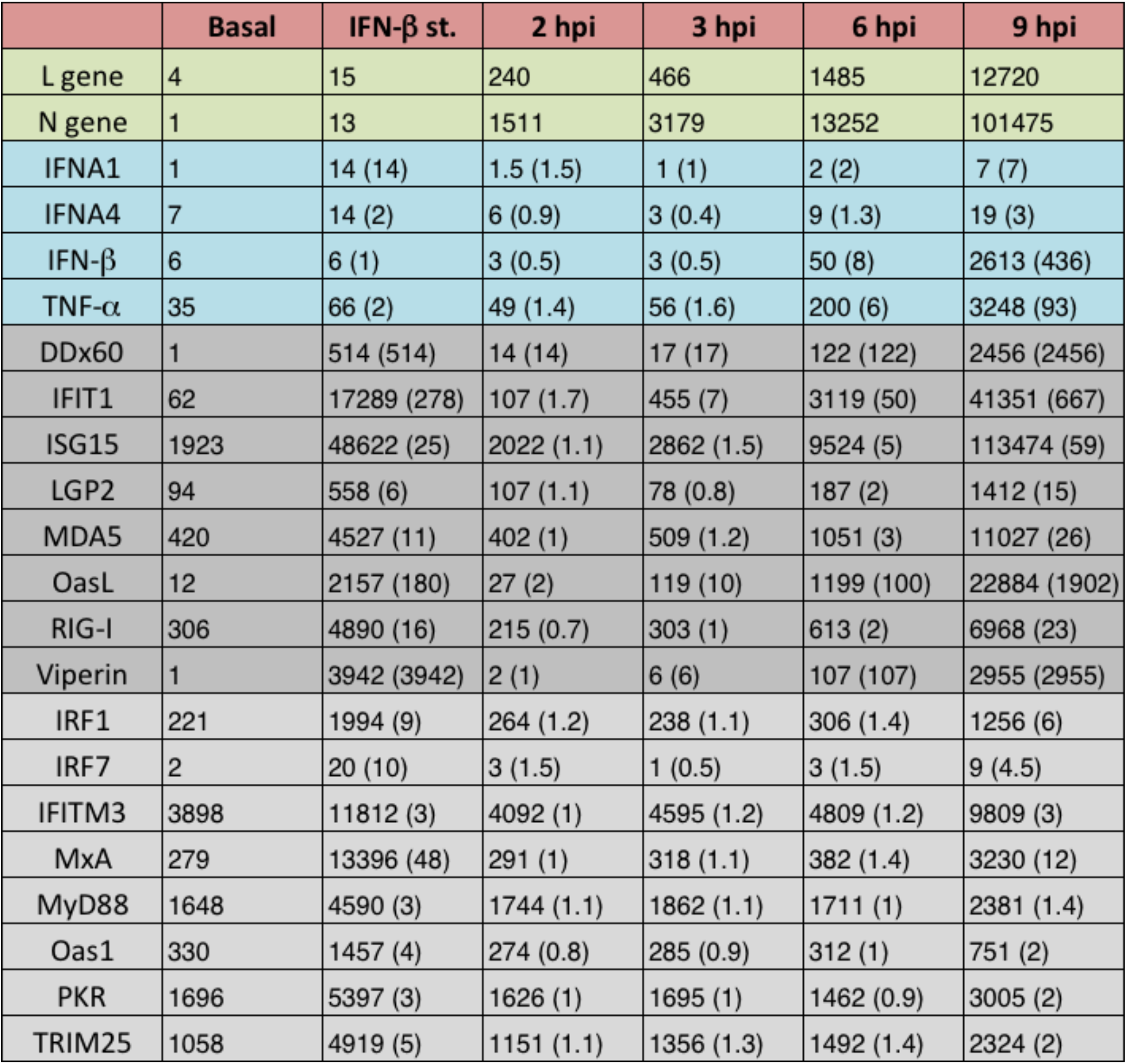
Number of transcripts detected by Nanostring nCounter gene expression system in control, IFN-β treated (2 hours), and SeV infected (2, 3, 6, and 9 hours) HepG2 cells. Number of transcripts is normalized to housekeeping gene GAPDH. Green rows denote viral genes, blue rows cytokines, dark gray rows early ISGs.

| | Basal | IFN-β st. | 2 hpi | 3 hpi | 6 hpi | 9 hpi |
| --- | --- | --- | --- | --- | --- | --- |
| L gene | 4 | 15 | 240 | 466 | 1485 | 12720 |
| N gene | 1 | 13 | 1511 | 3179 | 13252 | 101475 |
| IFNA1 | 1 | 14 (14) | 1.5 (1.5) | 1 (1) | 2 (2) | 7 (7) |
| IFNA4 | 7 | 14 (2) | 6 (0.9) | 3 (0.4) | 9 (1.3) | 19 (3) |
| IFN-β | 6 | 6 (1) | 3 (0.5) | 3 (0.5) | 50 (8) | 2613 (436) |
| TNF-α | 35 | 66 (2) | 49 (1.4) | 56 (1.6) | 200 (6) | 3248 (93) |
| DDx60 | 1 | 514 (514) | 14 (14) | 17 (17) | 122 (122) | 2456 (2456) |
| IFIT1 | 62 | 17289 (278) | 107 (1.7) | 455 (7) | 3119 (50) | 41351 (667) |
| ISG15 | 1923 | 48622 (25) | 2022 (1.1) | 2862 (1.5) | 9524 (5) | 113474 (59) |
| LGP2 | 94 | 558 (6) | 107 (1.1) | 78 (0.8) | 187 (2) | 1412 (15) |
| MDA5 | 420 | 4527 (11) | 402 (1) | 509 (1.2) | 1051 (3) | 11027 (26) |
| OasL | 12 | 2157 (180) | 27 (2) | 119 (10) | 1199 (100) | 22884 (1902) |
| RIG-I | 306 | 4890 (16) | 215 (0.7) | 303 (1) | 613 (2) | 6968 (23) |
| Viperin | 1 | 3942 (3942) | 2 (1) | 6 (6) | 107 (107) | 2955 (2955) |
| IRF1 | 221 | 1994 (9) | 264 (1.2) | 238 (1.1) | 306 (1.4) | 1256 (6) |
| IRF7 | 2 | 20 (10) | 3 (1.5) | 1 (0.5) | 3 (1.5) | 9 (4.5) |
| IFITM3 | 3898 | 11812 (3) | 4092 (1) | 4595 (1.2) | 4809 (1.2) | 9809 (3) |
| MxA | 279 | 13396 (48) | 291 (1) | 318 (1.1) | 382 (1.4) | 3230 (12) |
| MyD88 | 1648 | 4590 (3) | 1744 (1.1) | 1862 (1.1) | 1711 (1) | 2381 (1.4) |
| Oas1 | 330 | 1457 (4) | 274 (0.8) | 285 (0.9) | 312 (1) | 751 (2) |
| PKR | 1696 | 5397 (3) | 1626 (1) | 1695 (1) | 1462 (0.9) | 3005 (2) |
| TRIM25 | 1058 | 4919 (5) | 1151 (1.1) | 1356 (1.3) | 1492 (1.4) | 2324 (2) |

Up-regulation was also detected at 3 hpi for some of the early ISGs, such as *DDX60, IFIT1, OASL* and *Viperin*. A common property of these genes is that their basal level expression is very low, therefore when induced, they produce very high folds of increase with respect to the uninfected cells. On the other hand, induction of *ISG15, LGP2, MDA5* and *RIG-I* was not detectable at 3 hpi, because their basal expression levels were higher, therefore, induction in a small fraction of the cells did not produce a significant fold change when ensemble averaged (Table 1 & Table 2). On the other hand, smFISH data show simultaneous induction of *RIG-I, MDA5* and *LGP2* with bright transcription sites and at least 2-fold increase in the number of cytoplasmic transcripts (Table 1, Fig. 3A), in the same cells that activated transcription of *IFIT1*or *OASL* at 3 hpi, without exception. Overall, this comparison highlights the power of single cell approach with single molecule sensitivity to reveal gene expression patterns hidden in ensemble analysis.

In addition to their role in induction of *IFNB1*, transcription factors IRF3 and IRF7 have been shown to directly induce expression of some antiviral genes ^16, 34^. To test their roles in expression of the early ISGs, we immunofluorescently labeled IRF3 in SeV-infected cells, and analyzed the time-course of its nuclear translocation, which is required for its transcriptional activation role. We observed nuclear localization of IRF3 in 6 %, 25 % and 64% of the cells at 3, 6 and 9 hpi respectively, which is similar to that of *RIG-I* induction (Fig. 3*D*, *E,* Fig. 2*B*). Further, we used siRNA to knockdown IRF3 and/or IRF7. Following knockdown, we infected HepG2 cells for 9 hours with SeV, and performed qRT-PCR to measure the level of *RIG-I* mRNA. We found that SeV-induced expression of *RIG-I* is significantly reduced when both IRF3 and IRF7 are knockdown (Fig. 3*F*). These results together indicate that early ISGs expression is dependent on IRF3 or IRF7.

### Early ISGs expression may contribute to decision making process of *IFNB1* induction

smFISH analysis of *IFNB1* transcripts in single cells confirmed the heterogeneous nature of *IFNB1* expression among the cell population, which would otherwise be obscured in ensemble measurements. At any given time from 9 hpi onwards, we observed only a fraction of cells, typically less than 25%, expressing *IFNB1* as opposed to more than 70% showing *RIG-I*induction (Fig. 2*A*, *B*).

Several factors may shape the decision making process of *IFNB1* induction, one of which is the variation in the extent of viral replication ^20, 25^. We therefore performed smFISH analysis on SeV L gene mRNA which codes for the viral replicase to quantify viral replication. SeV-infected cells displayed viral mRNA while control cells did not show any signal above the background (Fig. 4*A*). At 9 hpi, we observed a high degree of cell-to-cell variation in L gene transcript counts, ranging from a few to several hundred regardless of whether *IFNB1* is expressed or not (Fig. 4B and Fig. S3) and determined a low Pearson’s correlation coefficient of only 0.26 between L gene and *IFNB1* mRNA counts (Fig. 4*B*), showing the amount of viral load is not a significant determinant of bimodal *IFNB1* expression in single cells.

**Figure 4.**
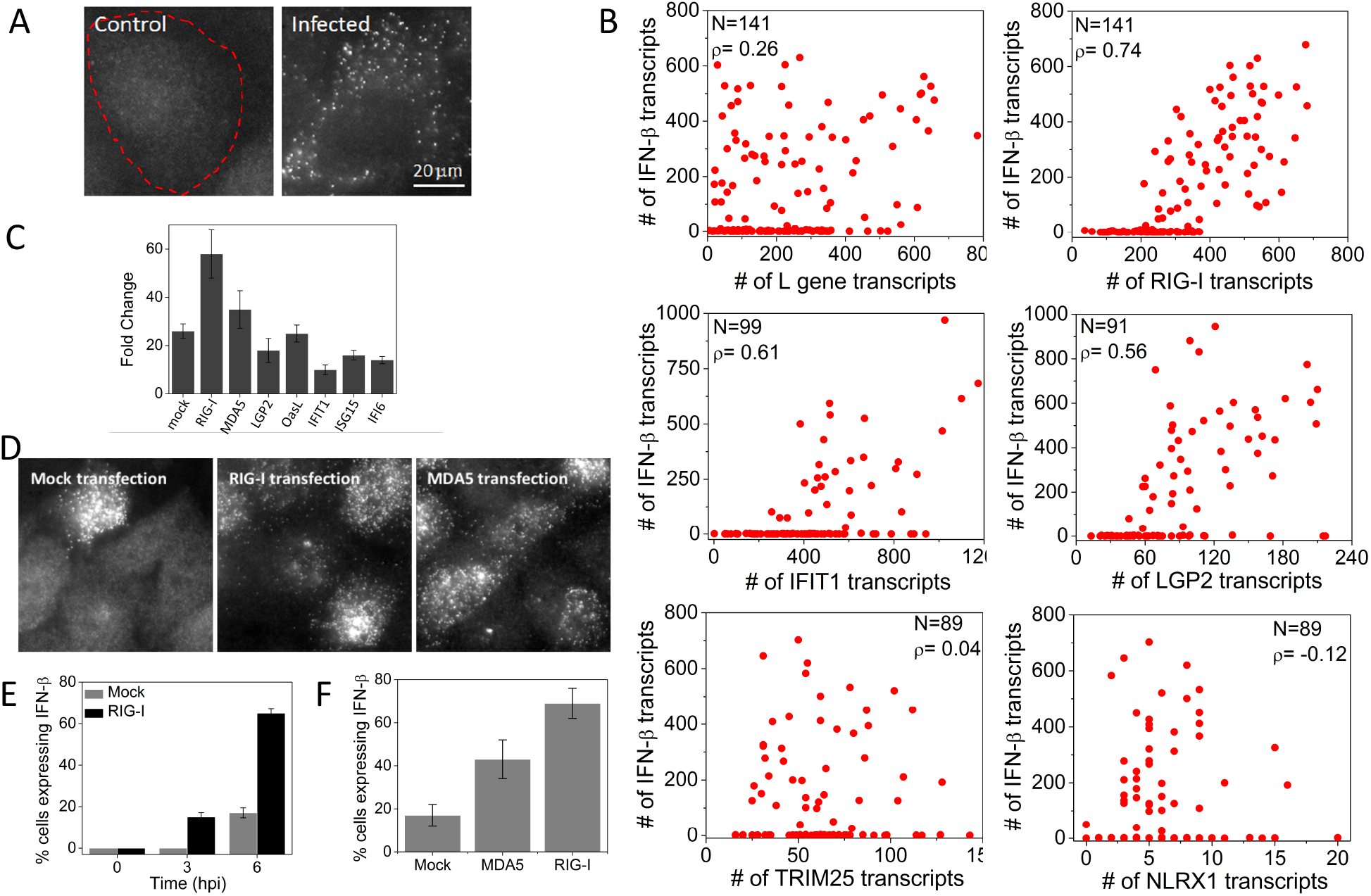
Early ISGs, but not viral L gene or other ISGs, are highly correlated with *IFNB1*expression. (A) Images of SeV L gene transcripts in control and SeV infected (6 hours) HepG2 cells. Cell border is denoted by dashed outline. (B) Scatter plots of number of L gene, *RIG-I*, *IFIT1*, *LGP2*, *TRIM25*, and *NLRX1* against *IFNB1* transcripts in single HepG2 cells at 9 hours post SeV infection. N: Number of cells analyzed, r: Pearson’s correlation coefficient. (C) qRT - PCR analysis of *IFNB1* mRNA in SeV infected (9 hours) HepG2 cells with prior plasmid expression (12 hours) of mock, *RIG-I*, *MDA5*, *LGP2*, *OASL*, *IFIT1*, *ISG15* and *IFI6*. Results are normalized to GAPDH, and presented as fold change relative to control cells. Error bars show standard deviation of two independent experiments. (D) Images of *IFNB1* transcripts in HeLa cells 6 h post SeV infection with prior transfection of an empty vector, *RIG-I* or *MDA5* (E) Percentage of HeLa cells (mean ± SD) expressing the *IFNB1* gene at different time points post SeV infection, with prior transfection of an empty vector or *RIG-I*. (F) Percentage of HeLa cells (mean ± SD) expressing the *IFNB1* gene at 6 hours post SeV infection, with prior transfection of an empty vector, *RIG-I* or *MDA5*.

In contrast to the L gene mRNA which was largely uncorrelated with *IFNB1* induction, we observed that every cell that expressed *IFNB1* also expressed *RIG-I*, as well as other early ISGs, including *MDA5, LGP2,* and *IFIT1*, over basal levels, and cells expressing higher amounts of early ISGs are more likely to be expressing *IFNB1* with high Pearson’s correlation coefficients (Fig. 4*B,* Fig. S4). On the other hand, ISGs that require IFN activity for induction such as *TRIM25* and *NLRX1* did not correlate with *IFNB1* induction (Fig. 4*B*).

The fact that RIG-I and MDA5 are major players in viral recognition and subsequent *IFNB1*induction, together with our data showing early *RIG-I* and *MDA5* expression precedes IFN, suggests that early expression of *RIG-I* and *MDA5* may have a causal role in inducing *IFNB1* in response to viral infection. In support of this hypothesis, we individually overexpressed *RIG-I, MDA5, LGP2, OASL, IFIT1, IFI6* and *ISG15* prior to viral infection by transfecting HepG2 cells with plasmids and monitored *IFNB1* expression following SeV infection by qRT-PCR. Transfection of *RIG-I* or *MDA5* encoding plasmids increased SeV-induced *IFNB1* production compared to mock transfected cells, but not other early ISGs, consistent with previous observations (Fig. 4*C*)^19, 22^. By smFISH analysis, we also showed that the percentage of *IFNB1*expressing HeLa cells significantly increased from about 20% to 60% at 6 hpi if the cells were transfected with *RIG-I* prior to viral infection. Additionally, we were able to detect *IFNB1* in about 18% of cells at 3 hpi (Fig. 4*D*, *E*). Likewise, transfection with MDA5 increased the percentage of *IFNB1* producing HeLa cells upon subsequent infection (Fig. 4*D*, *F*). Together, these experiments suggest that IFN-independent expression of *RIG-I* and *MDA5* may have a causal role in activating type I IFN in response to viral infection.

## Discussion

Although induction of ISGs through signaling triggered by type I IFN binding to the receptor is a major innate immune response, a more comprehensive picture is developing and showcases other mechanisms through which antiviral gene expression takes place ^16, 34^. In this study, using smFISH, we could identify a small fraction of cells showing up-regulation of RIG-I and several other genes upon viral infection, IFN-independently, whereas if we had relied on bulk measurements, such as qRT-PCR or Nanostring system, some of this information would have been lost due to the ensemble averaging over the entire population.

We exploited the multicolor nature of smFISH technology to study the well-known heterogeneity in the *IFNB1* expression in single cells^17-24^. Our results together suggest that amplification of *RIG-I* and *MDA5* in a fraction of cells at the early stages of infection may be a determinant of heterogeneous *IFNB1* induction. Our findings are in agreement with previous reports showing that *IFNB1* expressing cells have higher expression of RIG-I signaling pathway components ^22^ and heterogeneity in *IFNB1* expression is mainly of cellular origin ^23, 26^.

Dendritic cells are shown to rapidly sense and respond to pathogens by expressing antiviral genes, including *IFNB1*, by a few early responding cells and the cellular variation is then controlled by IFN mediated paracrine signaling from these early responding cells ^18, 26^. Patil et al have shown that secreted factors accounted for <10% of the total variation in single cell *IFNB1*responses, at the earliest time points *IFNB1* was detectable (40% of cells were expressing *IFNB1*). The fraction of cells expressing *IFNB1* then further increased over time due to paracrine signaling because treatment with drug Brefeldin A, which blocks paracrine signaling, blocked the increase in the fraction of cells expressing *IFNB1* ^24^. Because we mainly looked at the earliest time point *IFNB1* was detectable, the cell-to-cell variation is likely due to cell-intrinsic factors, not paracrine signaling.

The heterogeneity in *IFNB1* response has previously been shown to be relieved by increasing the concentration of various components of the pathway, such as RIG-I, TRIM25, IRF3, IRF7 and NF-kB to non-physiological levels ^19, 22^. In contrast, here we performed our smFISH experiments without disturbing the native system by forced expression of any of the components. We have found that *RIG-I*, but not *TRIM25, IRF3* or *IRF7*, is up-regulated in the native system prior to *IFNB1* induction, although over-expression of all of these components seemed to hyperactivate the *IFNB1* response in previous studies.

Further studies are required to understand the mechanism with which the early expression of *RIG-I* and *MDA5* facilitates induction of *IFNB1*. The obvious explanation is that the more RIG-I cells contain, the more responsive they are to viral infection. RIG-I and MDA5 have been shown to oligomerize upon viral infection, and the oligomeric forms of these proteins are highly active in inducing *IFNB1* ^35-37^. Amplification of RIG-I and MDA5 at the early stages of viral infection may promote oligomerization of these proteins, and subsequent aggregation of MAVS, and downstream signaling to *IFNB1* induction. RIG-I may also be promoting *IFNB1* expression by regulating NF-kB activity through binding to NF-kB 3’-UTR mRNA ^38^.

Finally, we still observe a fraction of *IFNB1* non-expressing cells showing high expression of early ISGs. Therefore, in addition to the expression of IRF3/7 dependent RIG-I and MDA5, there must be additional steps required for *IFNB1* induction, for example promoter binding and interchromosomal associations, providing further means of fine tuning the signaling process ^19^.

## Conclusion

Using a single-cell, single molecule technique, we could study the very early stages of antiviral gene expression and demonstrated a novelty in the expression kinetics of RLRs. Our results together suggest that there may be a link between the early expression of RIG-I and MDA5 and induction of the *IFNB1* gene.

## Materials and Methods

### Cell Culture, Virus Infection, IFN treatment, IFN neutralization and ELISA

HeLa and Vero cells were grown in DMEM supplemented with 10% FBS and 1% PENSTREP at 37°C. HepG2 cells were grown in DMEM supplemented with 10% FBS, 1% PENSTREP and 1mg/ml G418 at 37°C. Sendai Virus Cantell (SeV) was grown in 10 day old embryonated chicken eggs (Charles River Laboratories). Eggs were inoculated with 100 µl of undiluted virus in order to generate DI-rich virus stocks. Cells were seeded at least 24 hours prior to infection. On the day of infection, medium is replaced by fresh medium containing SeV at MOI 5. IFN-β treatments were carried out by adding 1000 units/ml of human IFN-β (PBL InterferonSource, 11410-2). Type I IFN activity was blocked by addition of 10 µg/ml of anti-IFN-α (Abcam, ab20200), 10 µg/ml anti-IFN-β (R&D systems, AF814), 10 µg/ml anti-IFN-λ (R&D Systems, MAB15981), and 5 µg/ml anti-IFNAR chain 2 (PBL InterferonSource, 21385-1) into the culture medium during infection. Verikine human IFN beta ELISA kit (PBL InterferonSource, 41410- 1B) was used to measure IFN-β from the culture medium.

### Plasmids, transfections and RNAi interference

Expression plasmids for RIG-I (pEF-flagRIG-Ifull) and MDA5 (pEF-flagMDA5full) were provided by Takashi Fujita.

A custom Gateway destination vector, pT-Rex™-DEST30/FLAG was created by using QuikChange mutagenesis to create a silent mutation NotI restriction site in the attR1 site in pT - Rex™-DEST30 (Invitrogen) by introducing C825A and G826A mutations using primers manufacturer-suggested primers (Stratagene QuikChange Lightning Kit). A dsDNA insert of the sequence

“gttgatgggcggccgctcgaaaacctgtattttcagggcactagtggcgacagcctgagctggctgctgagactgctgaacctgtgcacccccagccgggccgccctgctgaccggccggcctgcaggggactacaaagaccatgacggtgattataaagatcatgacatcgattacaaggatgacgatgacaagtagtaatgagggcccggagatg” was synthesized as a gBlok (IDT-DNA) and inserted via ligation into the NotI and AgeI sites of pT-Rex™-DEST30, producing a mammalian expression vector designed to append an amino acid sequence to the C-terminus of a hORF gene:

LPTFLYKVVDGRPLENLYFQGTSGDSLSWLLRLLNLCTPSRAALLTGRPAGDYKDHDGDYKDHDIDYKDDDDK-[STOP]

This sequence encodes a TEV protease recognition sequence, an SFP synthase tag, 13-aa aldehyde tag, and a 3x-FLAG-tag.

To obtain expression vectors pT-Rex™-DEST30/RIG-I/FLAG, pT-Rex™-DEST30/MDA5/FLAG, pT-Rex™-DEST30/LGP2/FLAG, pT-Rex™-DEST30/OasL/FLAG, pT-Rex™-DEST30/IFIT1/FLAG, pT-Rex™-DEST30/ISG15FLAG and pT-Rex™- DEST30/IFI6/FLAG, hORF plasmids from the CCSB Human ORFeome Collection (ThermoFisher) encoding *MDA5, LGP2, OasL, IFIT1, ISG15* and *IFI6* were cloned via Gateway directional LR cloning using LR Clonase II plus (Invitrogen) in 16 hours overnight reactions according to the manufacturer’s directions. LR reaction mixtures were transformed into chemically competent BL21-DH5-alpha cells and selected overnight on 100 ug/mL carbenicillin plates.

HeLa cells were transiently transfected with *RIG-I* (pEF-flagRIG-Ifull) and *MDA5* (pEF-flagMDA5full) using JetPrime^™^ transfection reagent (Polyplus transfection, France).

HepG2 cell were transiently reverse transfected with pT-Rex™-DEST30/RIG-I/FLAG, pT-Rex™-DEST30/MDA5/FLAG, pT-Rex™-DEST30/LGP2/FLAG, pT-Rex™-DEST30/OasL/FLAG, pT-Rex™-DEST30/IFIT1/FLAG, pT-Rex™-DEST30/ISG15FLAG and pT-Rex™-DEST30/IFI6/FLAG using JetPrime^™^ transfection reagent (Polyplus transfection, France).

Plasmid transfected cells were grown for 9 to 12 hours before viral infection. siRNAs targeting *IRF3* (sc-35710) and *IRF7* (sc-38011) were obtained from Santa Cruz Biotechnology. 30 pmol of each siRNA was transfected at a final concentration of ~45 nM, using Lipofectamine RNAiMAX (Invitrogen) transfection reagent. Cells transfected with siRNA were grown for 30 hours before viral infection.

### RNA extractions, qRT-PCR and nCounter

Total RNA was extracted using QIAShredder and RNeasy mini kit (Qiagen) according to the manufacturer’s instructions. RNA quality was determined by Agilent 2100 bioanalyzer, and only RNA with RIN above 9 was used for reverse transcription reactions. Total RNA (900 µg) was reverse transcribed using the iScript™ cDNA Synthesis Kit (BIO-RAD). qRT-PCR analysis was carried out using SYBR-Green (Applied Biosystems).

The primer sequences used for qRT-PCR were as follows:

*RIG-I* forward: GGTTTAGGGAGGAAGAGGTGC, reverse: AAGTGTGGCAGCCTCCATTG;

*MXA* forward: ATGCTACTGTGGCCCAGAAA, reverse: GGCGCACCTTCTCCTCATA;

*TNFA* forward: GAGTGACAAGCCTGTAGCC, reverse: GCTGGTTATCTCTCAGCTCCA;

*IFNB1* forward: GACGCCGCATTGACCATCTA, reverse: GTGACTGTACTCCTTGGCCT;

*GAPDH* forward: GACAGTCAGCCGCATCTTCT, reverse: AAATGAGCCCCAGCCTTCTC.

100 ng total RNA was hybridized to a custom gene expression CodeSet according to the NanoString Gene Expression Assay Manual and analyzed on an nCounter digital analyzer (NanoString Technologies). All data analysis was performed using the nSolver software analysis (freely available for download from NanoString Technologies). Briefly, counts are normalized for all target RNAs in all samples based on the positive control RNA to account for differences in hybridization efficiency and post-hybridization processing. Subsequently an mRNA content normalization is done using the housekeeping gene GAPDH.

### Immunofluorescence

Cells were fixed with 4% paraformaldehyde for 10 minutes and then permeabilized with 0.5% Triton-X 100 for 10 minutes. Cells were incubated with 3% BSA for 1 hour, and primary and secondary antibodies, each for 2 hours, at the recommended dilutions. Antibodies for IRF3 and NF-kB were obtained from Cell Signaling (11904P) and Abcam (ab119826) respectively. Whole procedure was performed at room temperature. Immunofluorescence images were obtained using Zeiss LSM 710 confocal scanning microscope equipped with a 40x oil-immersion objective (NA 1.3).

### Single molecule FISH

DNA oligos for FISH experiments are designed using software available online (http://www.singlemoleculefish.com/). FISH probes are ordered from Biosearch Technologies. For smFISH, cells were washed with PBS buffer, and fixed with 4% formaldehyde for 10 minutes at room temperature. After fixation, the cells were washed three times with PBS, before being permeabilized in 70% ethanol overnight. Cells were rehydrated in a solution of 10% formamide and 2x SSC for 5 minutes before hybridization. Hybridization reactions were carried out in 100 µl of hybridization buffer containing 10% dextran sulfate, 2 mM vanadyl-ribonucleoside complex, 0.02% RNAse-free BSA, 50 µg E.coli tRNA, 2x SSC, 10% formamide, and probe cocktail (5-50 ng) for overnight at 37°C. A coverslip was placed over the hybridization buffer to spread the small volume over the entire surface of the chamber, and also to prevent evaporation during the overnight incubation. After hybridization, the cells were washed twice for 30 minutes each at 37°C using a wash buffer containing 10% formamide and 2x SSC. During imaging, we used an oxygen scavenging mounting medium containing 10 mM Tris HCl pH 8.0, 2x SSC, 1% glucose, glucose oxidase (or pyranose oxidase) and catalase. We acquired smFISH images with a Zeiss Axiovert 200M inverted fluorescence microscope, equipped with a 100x oil-immersion objective (NA 1.45) and a Cascade 512b high sensitivity camera. Filter cubes to discriminate between Cy3, Alexa 594, and Cy5 were obtained from Semrock (SpOr-B-000-ZERO, SpRed-B-000-ZERO, and Cy5-4040C-000-ZERO respectively). 30 z-stacks were taken automatically with 0.3 microns between the z-slices, with an exposure time of 0.5 s.

Low magnification smFISH images (Supplementary Figure S2) images were obtained using Zeiss LSM 710 confocal scanning microscope equipped with a 40x oil-immersion objective (NA 1.3).

To identify and count the spots corresponding to individual mRNA molecules, we used a MATLAB program described previously ^27^. Briefly the program applies a Laplacian of Gaussian filter to the input TIFF images to enhance the spots. To identify spots, it applies an intensity threshold. The program counts number of spots detected for all possible threshold values in order to identify the appropriate threshold. When the number of spots identified is plotted as a function of the threshold value, a plateau region is observed, where the spot count does not vary significantly over a broad range of thresholds. The threshold is manually selected at the plateau region, based on the graph and visual feedback.

## Acknowledgements

We would like to thank Mayandi Sivaguru and Glenn Fried (microscopy), Benjamin J. Leslie (cloning) for expert help. We would like to thank Arjun Raj and Kaushik Ragunathan for helpful discussions. This research was partly funded by NIAID grants U19 AI083025 (to TH, SM and AG-S), by NIAID grant U19AI117873 (to AG-S).

